# Learning interpretable structural similarity from tandem mass spectra for small molecule analog discovery

**DOI:** 10.64898/2026.06.17.733050

**Authors:** Juan Sebastian Piedrahita Giraldo, Katyeny Manuela Da Silva, Mohammad Reza Zare Shahneh, Mingxun Wang, Kris Laukens, Thomas De Vijlder, Wout Bittremieux

## Abstract

Analog discovery remains a central bottleneck in mass spectrometry-based untargeted metabolomics, as conventional spectral similarity scores poorly reflect molecular structure. We introduce SIMBA, a transformer-based model that infers two interpretable graph-based distances, maximum common edge subgraph and substructure edit distance, directly from tandem mass spectra. SIMBA consistently retrieves structurally closer analogs than existing methods, enabling structure-aware small molecule identification beyond exact spectral matching.

---

The identification of small molecules underpins a wide range of scientific and translational applications, such as in metabolomics, natural products research, and exposomics. In these fields, mass spectrometry (MS) is the primary analytical technology for molecular characterization, due to its sensitivity, throughput, and broad chemical coverage. Traditionally, tandem mass spectrometry (MS/MS)-based identification relies on exact spectral matching, in which experimental spectra are compared against reference spectra in curated libraries [1]. While effective when reference spectra are available, this approach inherently limits identification to previously characterized compounds.

In practice, many real-world analyses aim to identify structural analogs rather than exact molecular matches—molecules that share substantial substructural similarity but differ by specific functional groups or scaffolds. This scenario is common in untargeted studies, where the chemical diversity encountered in biological and environmental samples far exceeds the coverage of existing spectral libraries. Analog discovery is therefore substantially more challenging, as structurally related molecules often lack direct spectral references. Although MS/MS spectra encode rich structural information, widely used similarity measures such as the modified cosine score frequently fail to reflect true structural relatedness when fragmentation patterns differ [2, 3]. These methods rely primarily on peak alignment, which can capture only limited structural changes, often corresponding to a single modification, and break down when analogs exhibit divergent fragmentation behavior.

To address these limitations, several machine learning approaches have been proposed to learn spectral representations tailored to analog searching. Spec2Vec [3] interprets mass spectra as documents and applies the Word2Vec algorithm to learn embeddings based on co-occurring fragment ions. MS2DeepScore [4] instead employs a twin neural network trained on pairs of spectra annotated with molecular fingerprint-based Tanimoto similarity scores. DreaMS [5] is an MS/MS foundation model, which can be finetuned for exact and analog searching. While these approaches improve over traditional alignment-based metrics, a major limitation is that they rely on fingerprint-derived similarity measures that are difficult to interpret in terms of concrete chemical transformations. Furthermore, a disadvantage of earlier approaches is that they use neural network architectures with limited representational capacity, which restricts their ability to capture long-range dependencies and complex non-linear relationships between molecular structure and fragmentation behavior.

A fundamental choice for analog discovery is how to represent structural similarity. The Tanimoto similarity is the most widely used measure in cheminformatics [6], but its reliance on binary fingerprints can obscure fine-grained structural differences and lead to substantial information loss [7]. Graph-based similarity measures provide a more chemically grounded alternative. In particular, the maximum common edge subgraph (MCES) distance measures structural dissimilarity as the number of chemical bonds that differ between two molecular graphs [8], providing a direct and interpretable bond-level description of molecular divergence.

Here, we further introduce the “substructure edit distance,” a complementary metric that quantifies the number of bond-level attachments or removals required to transform two molecules into their maximum common substructure. Unlike fingerprint-based similarities, this metric explicitly captures discrete structural modifications and aligns conceptually with the fragment-based nature of MS/MS spectra, which reflect the presence or absence of molecular substructures. Together, MCES distance and substructure edit distance provide interpretable, chemically meaningful targets for learning structural similarity directly from spectral data.

Building on these metrics, we developed SIMBA (Spectral Inference of Molecular Bio-Analogs), a transformer-based neural network that infers both distances directly from spectral data to enable accurate analog discovery (Figure 1a). These two metrics provide complementary views of molecular relatedness: the substructure edit distance captures the number of discrete structural modifications between molecules, whereas the MCES distance quantifies the total number of differing chemical bonds underlying those modifications.

**Figure 1:**
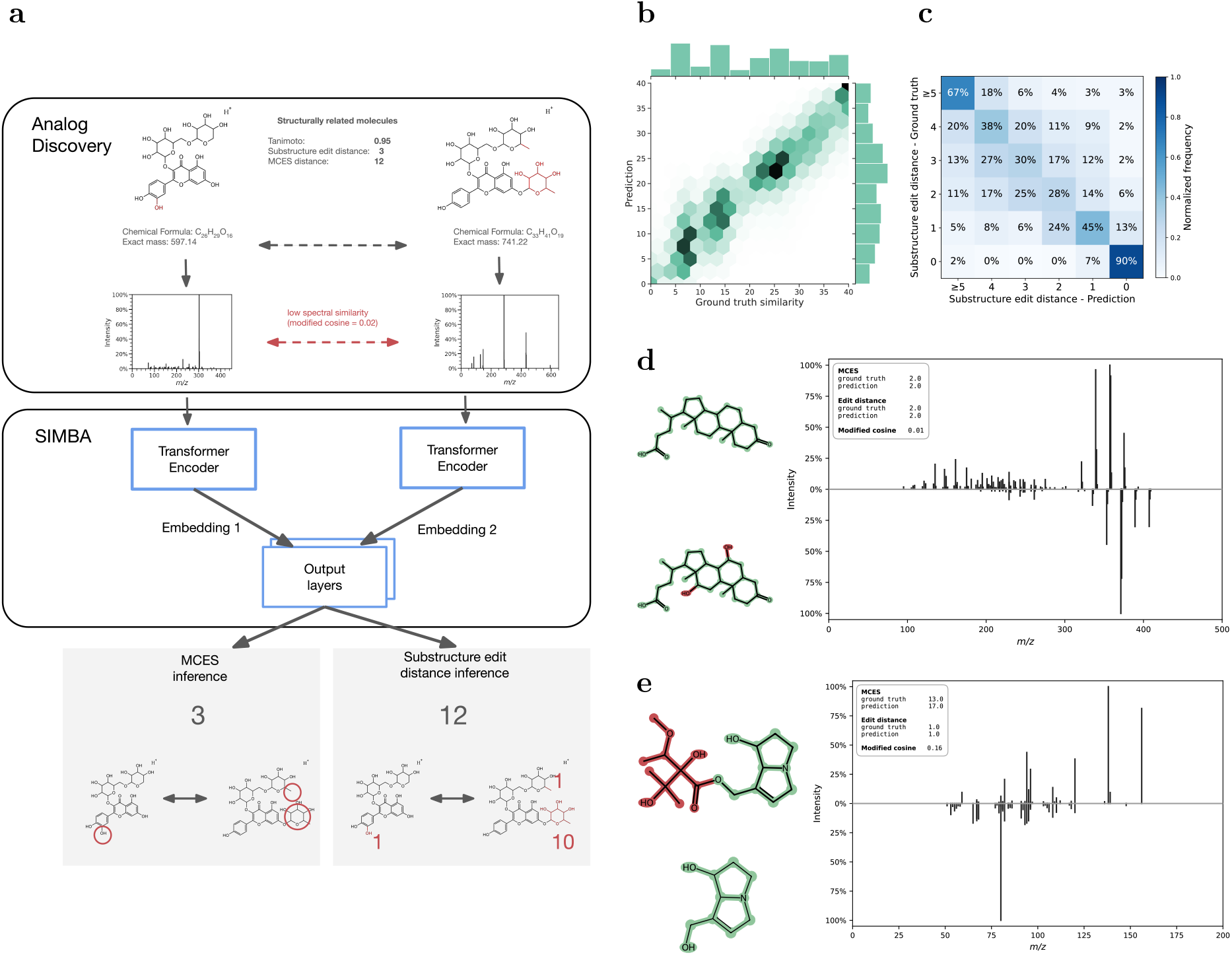
SIMBA learns chemically meaningful structural similarity from MS/MS spectra. **(a)** Structural similarity between molecules is not necessarily reflected by conventional spectral similarity, indicating the need for learned, structure-aware representations. We propose SIMBA, a twin transformer-based encoder model that infers substructure edit distance and MCES distance directly from MS/MS spectra. **(b)** Inferred MCES distances strongly correlate with ground truth values across the test set (Spearman *r* = 0.93), demonstrating accurate recovery of bond-level structural differences from spectral data. **(c)** Substructure edit distance is inferred with high fidelity, achieving 50 % accuracy on a balanced six-class ordinal classification task. **(d, e)** Representative examples illustrating SIMBA’s ability to jointly capture discrete structural edits and overall bond differences. Shared substructures are highlighted in green and differing substructures in red. **(d)** Bile acids 3-ketolithocholic acid and 3-oxocholic acid, differing by two structural edits (edit distance 2; MCES distance 2), both correctly inferred. **(e)** Alkaloids europine and heliotridine differing by a single substructure modification (edit distance 1; MCES distance 13), with accurate edit distance inference and a near-exact MCES estimate.

SIMBA was trained on more than 200 million molecular pairs derived from the NIST20 and MassSpecGym [9] spectral libraries, with compounds partitioned into disjoint training, validation, and test sets based on Murcko scaffolds (205 million, 3.7 million, and 3.6 million pairs, respectively). On the test set, SIMBA achieved a Spearman correlation of 0.93 between inferred and ground truth MCES distances, indicating strong agreement across a wide range of structural dissimilarities (Figure 1b, Supplementary Figure 1a). For substructure edit distance inference, SIMBA obtained a multi-class accuracy of 50 % on a balanced six-class ordinal classification task (edit distances 0–4 and ≥ 5; Figure 1c, Supplementary Figure 1b). Given the ordinal nature of the problem and the increasing ambiguity associated with larger structural differences, this performance demonstrates that SIMBA can reliably infer chemically meaningful edit distances directly from spectral data. Representative examples illustrate that both inferred MCES and substructure edit distances closely track the true structural differences between molecular pairs, highlighting SIMBA’s ability to capture fine-grained variations in molecular structure from MS/MS fragmentation patterns (Figure 1d,e).

We next evaluated SIMBA in an analog discovery setting using query spectra from the CASMI 2022 dataset, searching against two reference libraries: a combined NIST20 + MassSpecGym library and the GNPS community libraries [10]. SIMBA and DreaMS, both transformer-based methods, perform similarly in terms of median normalized MCES for the top-ranked analog (Figure 2a, Supplementary Figure 2a). However, SIMBA exhibits fewer extreme outlier inferences and, more importantly, substantially outperforms DreaMS in an analog classification setting (Figure 2b, Supplementary Figure 2b). Furthermore, both SIMBA and DreaMS clearly outperform alignment-based and other deep learning approaches in terms of overall ranking performance (Figure 2c, Supplementary Figure 2c), highlighting SIMBA’s ability to robustly discriminate true analogs. Notably, this strong performance persisted even when candidate ranking was evaluated using Tanimoto similarity rather than MCES distance (Supplementary Figure 3), indicating that the learned spectral representations generalize beyond the specific structural metrics used during training.

**Figure 2:**
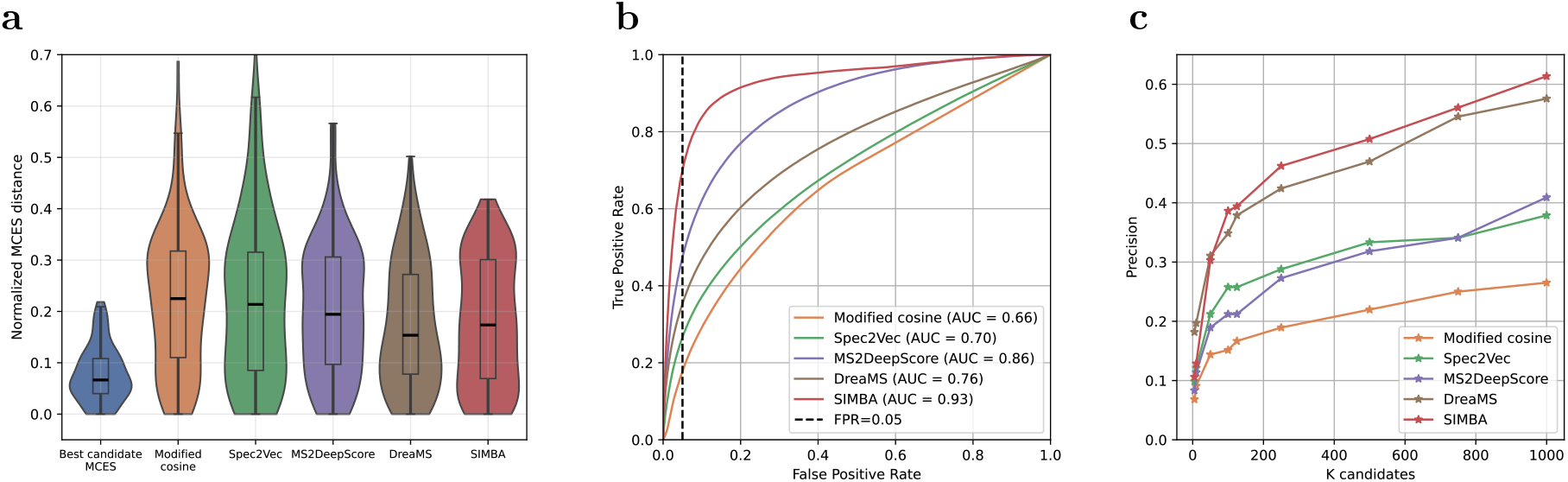
SIMBA outperforms existing methods in analog discovery. **(a)** Analog search performance using CASMI 2022 query spectra against the NIST20 + MassSpecGym reference library. Structural similarity of the top-ranked analog is quantified using the normalized MCES distance (MCES rescaled by molecular size; lower values indicate greater structural similarity). The central line indicates the median, the box bounds the interquartile range (25th–75th percentiles), and the whiskers indicate 1.5 times the interquartile range. SIMBA achieves comparable median normalized MCES (0.17) to DreaMS (0.15) while exhibiting fewer outliers, and consistently outperforms modified cosine (0.23), Spec2Vec (0.21), and MS2DeepScore (0.19). **(b)** Receiver operating characteristic (ROC) curves for discriminating true analogs from unrelated molecules, where pairs are binarized using a normalized MCES distance threshold of 0.3. At a false positive rate of 5 %, SIMBA achieves substantially higher true positive rates than competing approaches (SIMBA: 70 %; DreaMS: 36 %; MS2DeepScore: 47 %; Spec2Vec: 25 %; modified cosine: 18 %). **(c)** Ranking performance for analog retrieval. SIMBA consistently places the closest analog among the top candidates, performing similarly to DreaMS and outperforming other baselines.

In summary, SIMBA introduces a structure-aware approach for analog discovery that learns chemically interpretable similarity directly from MS/MS spectra by inferring complementary graph-based distances. This enables more reliable retrieval and ranking of structural analogs in untargeted analyses, supporting more accurate small molecule annotation across metabolomics, natural products research, and environmental chemistry.

## Methods

### Structural similarity metrics

SIMBA infers two complementary structural similarity metrics directly from MS/MS spectra: the maximum common edge subgraph (MCES) distance and the substructure edit distance.

The MCES distance quantifies structural dissimilarity between molecules represented as graphs, *G*_1_ = (*V*_1_, *E*_1_) and *G*_2_ = (*V*_2_, *E*_2_), where *V* denotes atoms and *E* denotes bonds. It is defined as:

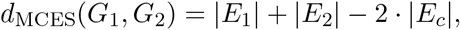

where |*E*_*c*_| is the number of shared edges in the maximum common edge subgraph of *G*_1_ and *G*_2_. Intuitively, this distance corresponds to the minimum number of bond modifications required to make the molecular graphs isomorphic [8].

Computing MCES is NP-hard; in this study, we employ the myopic MCES implementation (version 1.0.1) [8], which balances computational efficiency and accuracy. A distance threshold of 20 was applied: exact MCES values were computed below the threshold, while approximate values were returned for larger distances.

The substructure edit distance captures the number of discrete bond-level modifications needed to convert one molecule into another via their maximum common substructure (MCS). For molecules represented as graphs *G*_1_ = (*V*_1_, *E*_1_) and *G*_2_ = (*V*_2_, *E*_2_), let:

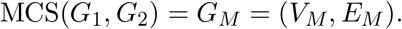

Attachment edges are defined as:

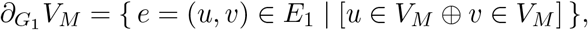

and 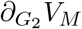 is defined analogously, where ⊕ denotes the exclusive OR. These edges correspond to bonds diverging from the shared core. The substructure edit distance is then:

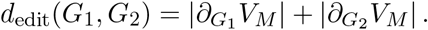

To ensure structural relevance and computational tractability, molecular pairs were filtered as follows: (i) Tanimoto similarity (2048-bit RDKit fingerprints) below 0.2 were excluded, (ii) molecules containing more than 60 heavy atoms (C, O, N, P, S, halogens) were removed, and (iii) pairs with an MCS containing fewer than 50% of the atoms of either molecule were discarded. Only pairs passing all filters were processed with RDKit (version 2023.9.4) [11] to determine the MCS and compute the corresponding substructure edit distance.

### Training data

SIMBA was trained on MS/MS spectra from the NIST20 and MassSpecGym [9] libraries, comprising 1,026,712 and 231,104 spectra, respectively. Following preprocessing to remove low-quality spectra and retain only singly protonated ions as precursors, 328,555 spectra corresponding to 42,774 unique compounds were included.

Preprocessing consisted of several steps to ensure data quality and consistency: (i) spectra lacking valid InChI or SMILES annotations were excluded; (ii) only spectra with singly charged, protonated precursor ions were retained; (iii) fragment peaks with intensity lower than 1% of the base peak intensity were removed and if applicable the number of fragments per spectrum was further limited to the 100 most intense peaks; (iv) peaks within ±0.1 *m*/*z* of the precursor ion were removed to eliminate residual precursor signal; (v) spectra containing fewer than 6 peaks were discarded; (vi) and fragment intensities were square-root transformed to enhance the influence of low-intensity peaks, followed by L2 normalization to standardize spectral intensity scales across spectra.

To ensure chemical diversity and prevent overfitting, compounds were partitioned based on their Murcko scaffolds [12], which capture the core structural framework of each molecule. Scaffolds were assigned to training (80%), validation (10%), and test (10%) sets, resulting in 262,316, 34,491, and 31,748 spectra per set, respectively. This scaffold-based partitioning guarantees that structurally distinct chemical classes are separated across splits, supporting robust evaluation of model generalization.

Next, pairwise structural similarity labels were computed for all compound pairs within each split. This resulted in 205,114,315 unique molecular pairs for training, 3,716,098 unique molecular pairs for validation, and 3,670,290 unique molecular pairs for testing, with both MCES and substructure edit distances calculated for each pair (Figure 3).

**Figure 3:**
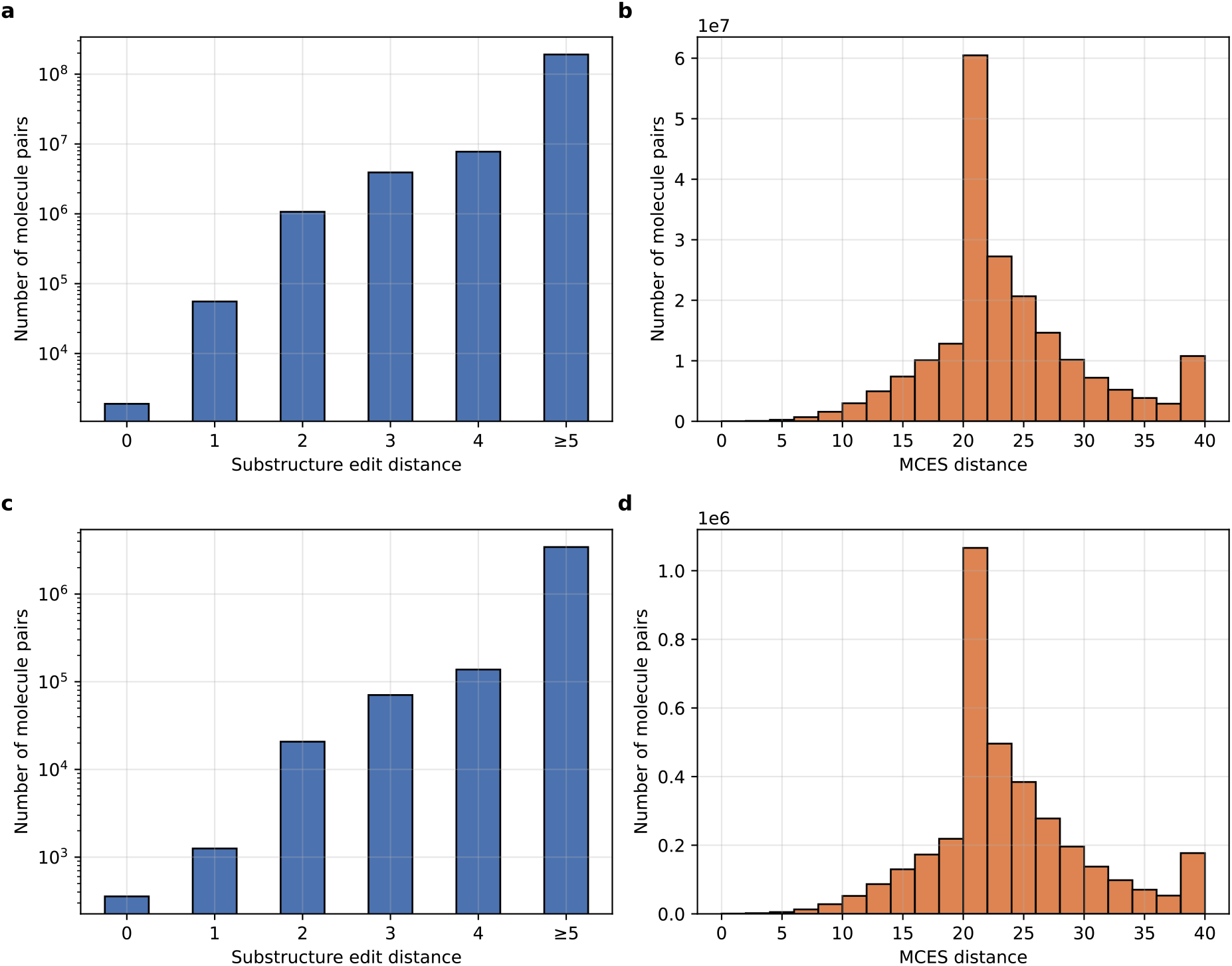
Training data characteristics. Distribution of substructure edit distance and MCES distance for the training set (**a, b**) and test set (**c, d**). The peak at MCES = 20 in panels **b** and **d** arises from the use of the myopic MCES approximation, where distances above this threshold are approximated. MCES values were capped at a maximum of 40.

### Neural network architecture

SIMBA is a twin transformer-based neural network designed to jointly infer (i) the MCES distance and (ii) the substructure edit distance between pairs of MS/MS spectra (Figure 4).

**Figure 4:**
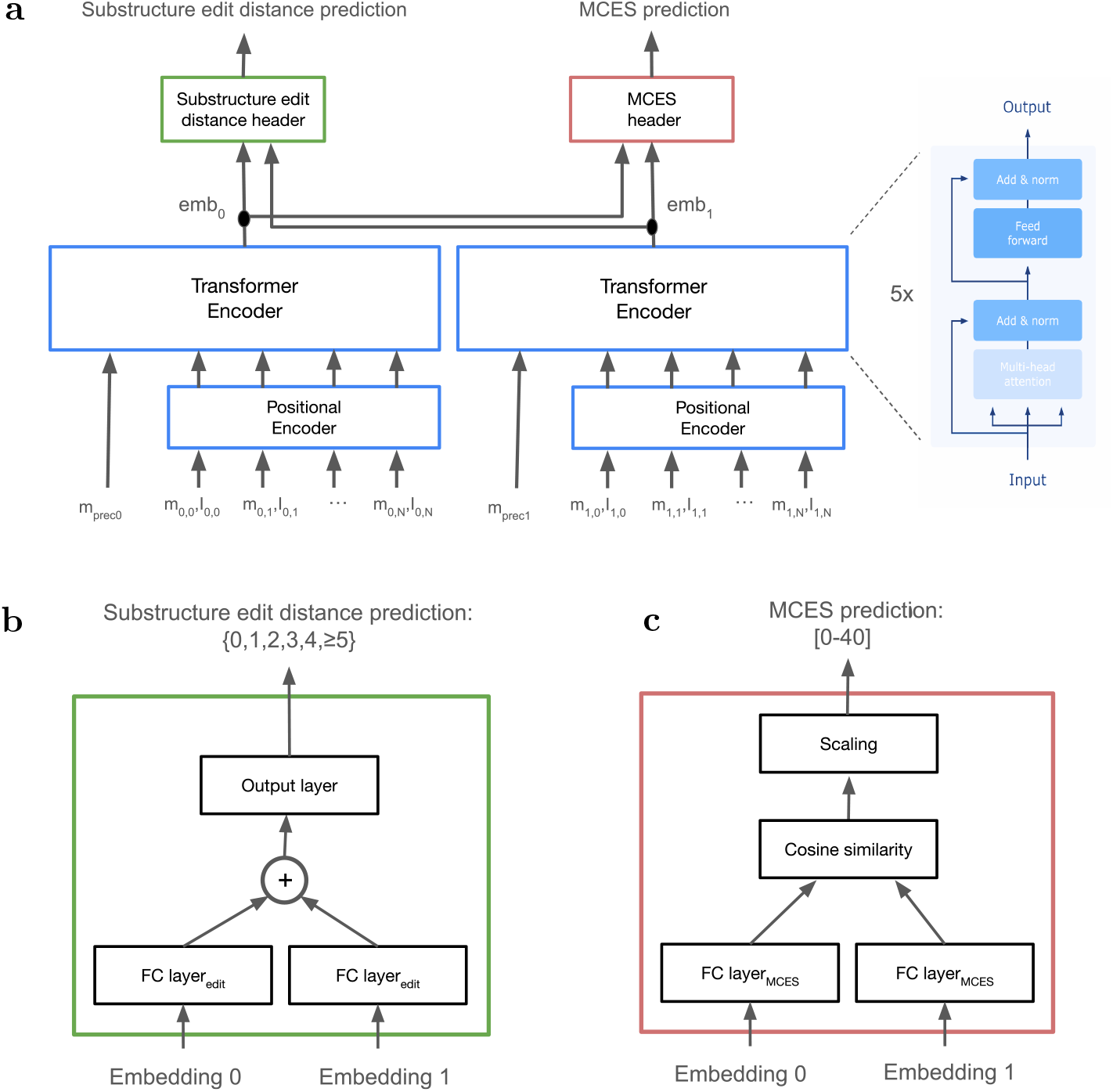
Architecture of SIMBA. **(a)** Overview of the twin transformer framework: two weight-shared spectrum encoders independently map input MS/MS spectra to latent embeddings, which are subsequently used for joint inference of structural similarity. **(b, c)** Task-specific inference heads. **(b)** For substructure edit distance, transformed embeddings are combined by summation and passed to a classification layer inferring ordinal classes. **(c)** For MCES distance, transformed embeddings are compared using cosine similarity, followed by linear rescaling to obtain a continuous distance inference.

The model consists of two identical branches with shared weights, each encoding one input spectrum into a latent representation. These embeddings are subsequently combined through task-specific inference heads.

Each input spectrum is represented as a set of up to 100 fragment peaks, defined by their *m*/*z* values and intensities, together with the precursor *m*/*z*. Spectral peaks are embedded using sinusoidal positional encodings applied separately to the *m*/*z* and intensity dimensions (wavelengths from 0.001 to 10,000 *m*/*z* for the *m*/*z* values, and from 10^*−*6^ to 1 for the intensities), enabling the model to capture both mass relationships and intensity patterns. The precursor *m*/*z* is prepended as an additional token to the sequence.

Each branch employs a spectrum transformer encoder [13] (implemented using the Depthcharge library, version 0.3.2) consisting of five stacked transformer blocks with a hidden dimension of 256. The final spectrum representation is obtained from the encoder output corresponding to the prepended precursor token, yielding a fixed-dimensional embedding vector of size 256.

The two spectrum embeddings are used to jointly infer structural similarity via two task-specific heads. For substructure edit distance inference, each embedding is first transformed through a fully connected layer (256 units). The resulting vectors are summed and passed through a final linear layer producing a six-dimensional output corresponding to the ordinal classes {0, 1, 2, 3, 4, ≥ 5}.

For MCES inference, each embedding is independently transformed through a fully connected layer (256 units), after which cosine similarity between the resulting vectors is computed. The similarity score is linearly rescaled to the range 0–40 to match the distribution of MCES distances in the training data. Empirically, this approach to model MCES via cosine similarity between learned embeddings yielded improved performance compared to direct regression using fully connected layers.

### Loss functions and training strategy

The distributions of both MCES and substructure edit distances are highly imbalanced, with a strong skew toward low-similarity pairs (Figure 3). To mitigate this imbalance and ensure adequate representation of structurally similar pairs, we employed distance-based weighting and resampling strategies during training.

Inference of the substructure edit distance was formulated as an ordinal classification task with six classes {0, 1, 2, 3, 4, ≥ 5}. To address class imbalance, molecular pairs were grouped into these bins and sampled with probabilities inversely proportional to their frequency:

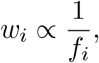

where *f*_*i*_ denotes the number of pairs in bin *i*. During training, a random spectrum was sampled on-the-fly for each compound pair to increase data diversity.

To account for the ordinal nature of the substructure edit distance, we used a cross-entropy loss with a distance-aware soft target matrix *M* ∈ ℝ^*C×C*^, in which target mass is concentrated on the true class and decays quadratically with ordinal distance:

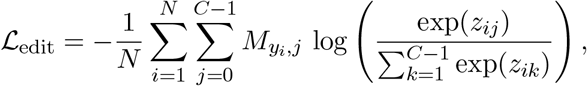

where *z*_*ij*_ denotes the logit for sample *i* and class *j*, and *y*_*i*_ ∈ {0, …, *C* − 1} is the ground-truth class label.

The distance-aware soft target matrix is defined as:

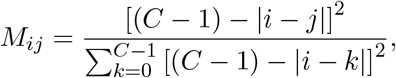

with *C* = 6. Rows are normalized and globally rescaled such that max_*i,j*_ *M*_*ij*_ = 1. This construction yields a symmetric matrix whose entries decrease quadratically with ordinal distance from the true class, thereby assigning greater target weight to near-miss classes than to more distant misclassifications.

MCES inference was formulated as a regression task. To emphasize accurate inference of small distances, which are most relevant for analog discovery, we applied a logarithmically derived nonlinear transformation to a rescaled version of the MCES distance. The loss is defined as:

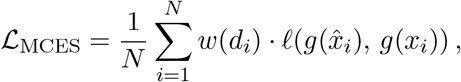

where *ℓ*(·, ·) denotes the mean squared error, *w*(*d*_*i*_) is a distance-dependent weight computed from the original MCES distance, and *g*(·) is defined as:

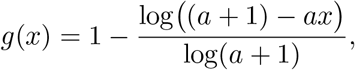

with *a* = 1 in our experiments.

Here, *x* denotes a scaled variable derived from the MCES distance:

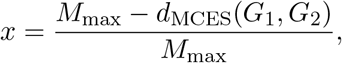

where *M*_max_ represents the maximum MCES distance (set to 40).

Under this formulation, identical structures (*d*_MCES_(*G*_1_, *G*_2_) = 0) map to *x* = 1, while maximally dissimilar structures map to *x* = 0. The nonlinear transformation *g*(·) increases sensitivity to differences among structurally similar molecules, thereby emphasizing accurate inference of small MCES distances, while avoiding numerical instabilities associated with direct logarithmic transformations.

To counteract imbalance, MCES distances in the range 0–40 were partitioned into 10 bins, and weights were assigned inversely proportional to bin frequency using the same scheme as for the substructure edit distance. This encourages the model to focus on underrepresented low-distance (high-similarity) pairs.

The final training objective combines both tasks using uncertainty-based weighting [14]:

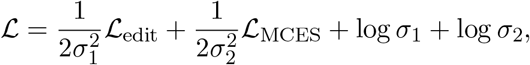

where *σ*_1_ and *σ*_2_ are learnable parameters. This formulation dynamically balances the contributions of the classification and regression objectives, improving training stability and preventing either task from dominating optimization.

### Model training

To improve robustness to experimental variability and enhance generalization, data augmentations were applied probabilistically and independently to each input MS/MS spectrum during training. Specifically, precursor mass values were perturbed by adding uniform noise up to 1% of the precursor *m*/*z*, simulating instrument measurement variability and reducing reliance on precursor mass alone. Additionally, peak dropout was applied by randomly removing up to 10% of fragment peaks per spectrum, mimicking incomplete fragmentation patterns and encouraging the model to learn robust representations from partial spectral information.

The primary SIMBA model was trained on the combined NIST20 and MassSpecGym datasets to maximize chemical diversity and coverage. To support reproducibility and open use, an additional model was trained exclusively on 70,135,880 pairs derived solely from MassSpecGym, excluding proprietary NIST20 data. Despite the reduced training set size, the public model achieved comparable performance to the full model (Supplementary Figure 4), demonstrating that SIMBA can learn meaningful structure–spectrum relationships from public data alone.

### Analog discovery

Model performance for analog discovery was evaluated using the CASMI 2022 dataset as query spectra. Two reference libraries were considered: (i) a combined NIST20 and MassSpecGym library, and (ii) the GNPS community spectral libraries [10]. The same preprocessing pipeline described above was applied to all library spectra, resulting in 53,618 GNPS spectra retained for searching.

The CASMI 2022 dataset consists of 158 MS/MS spectra from structurally diverse compounds. Spectra whose corresponding compounds were present in the reference libraries were excluded, yielding 132 query spectra.

For each query spectrum, SIMBA was used to infer MCES and substructure edit distances to all candidate library spectra. Candidates were ranked primarily by inferred MCES distance, reflecting its higher granularity as a structural similarity metric. Ties in MCES inferences were resolved using the substructure edit distance.

For each query, the top 10 candidates with the lowest inferred MCES distances were selected, from which the best candidate (lowest true MCES distance) was selected as the retrieved analog, following the same procedure as proposed by Huber et al. [3]. Performance was compared against baseline methods, including modified cosine similarity, Spec2Vec (version 0.8.0), MS2DeepScore (version 2.5.0), and DreaMS (version 1.0.0), using identical candidate ranking procedures.

To enable comparison across compounds of different sizes, retrieval quality was assessed using the normalized MCES distance between each query and its selected match, defined as:

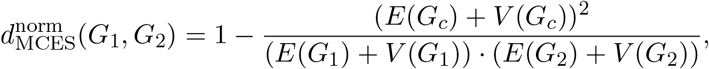

The fraction represents the Johnson similarity between the two molecules [15], with *G*_*c*_ the maximum common edge subgraph of molecules *G*_1_ and *G*_2_, and *E*(·) and *V* (·) the number of edges and vertices of respective molecules.

Ranking performance was further evaluated using recall@k, defined as the fraction of queries for which the best structural analog is found within the top *k* candidates.

## Data availability

Model weights for the SIMBA version trained using only public MS/MS spectra from MassSpecGym, the corresponding training data, and the CASMI 2022 query data can be found at https://zenodo.org/records/15275257.

Universal spectrum identifiers [16] for spectra visualized in Figure 1c,d are:

- 3-ketolithocholic acid: `mzspec:GNPS:BILELIB19:accession:CCMSLIB00005464754`
- 3-oxocholic acid: `mzspec:GNPS:BILELIB19:accession:CCMSLIB00005464724`
- europine: `mzspec:GNPS:PYRROLIZIDINE-ALKALOID-SPECTRAL-LIBRARY:accession: CCMSLIB00014205810`
- heliotridine: `mzspec:GNPS:PYRROLIZIDINE-ALKALOID-SPECTRAL-LIBRARY:accession: CCMSLIB00014205818`

## Code availability

All code for training and running SIMBA, as well as additional documentation, is available as open source under the Apache 2.0 license at https://github.com/bittremieuxlab/simba. Results presented in this manuscript were obtained using SIMBA version 1.0.

## Acknowledgements

This work was supported in part by the Research Foundation – Flanders (FWO 12A4025N and G0AHY25N).

## Author contributions

Juan Sebastian Piedrahita Giraldo: conceptualization, funding acquisition, investigation, methodology, software, validation, visualization, writing – original draft, writing – review & editing. Katyeny Manuela Da Silva: data curation, investigation, writing – review & editing. Mohammad Reza Zare Shahneh: methodology, software, writing – review & editing. Mingxun Wang: conceptualization, funding acquisition, methodology, supervision, writing – review & editing. Kris Laukens: conceptualization, funding acquisition, supervision, writing – review & editing. Thomas De Vijlder: conceptualization, funding acquisition, supervision, writing – review & editing. Wout Bittremieux: conceptualization, funding acquisition, methodology, project admninistration, resources, supervision, writing – original draft, writing review & editing.

## Supporting information

**Supplementary Figure 1:**
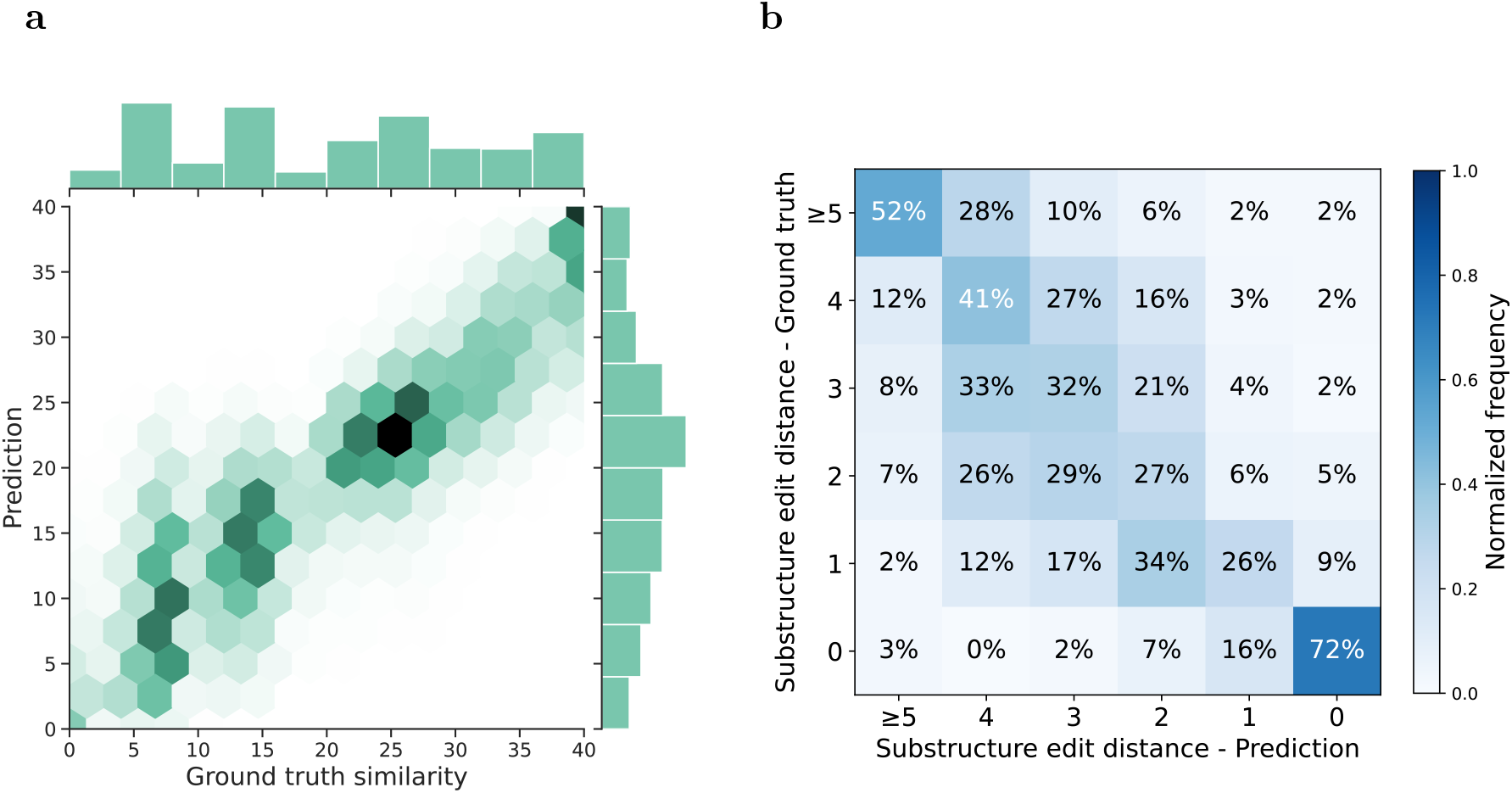
Test set performance of SIMBA trained on public MassSpecGym data. **(a)** Inferred MCES distances strongly correlate with ground truth values (Spearman *r* = 0.89). **(b)** Substructure edit distance is inferred with high fidelity, achieving 42 % accuracy on a balanced six-class ordinal classification task.

**Supplementary Figure 2:**
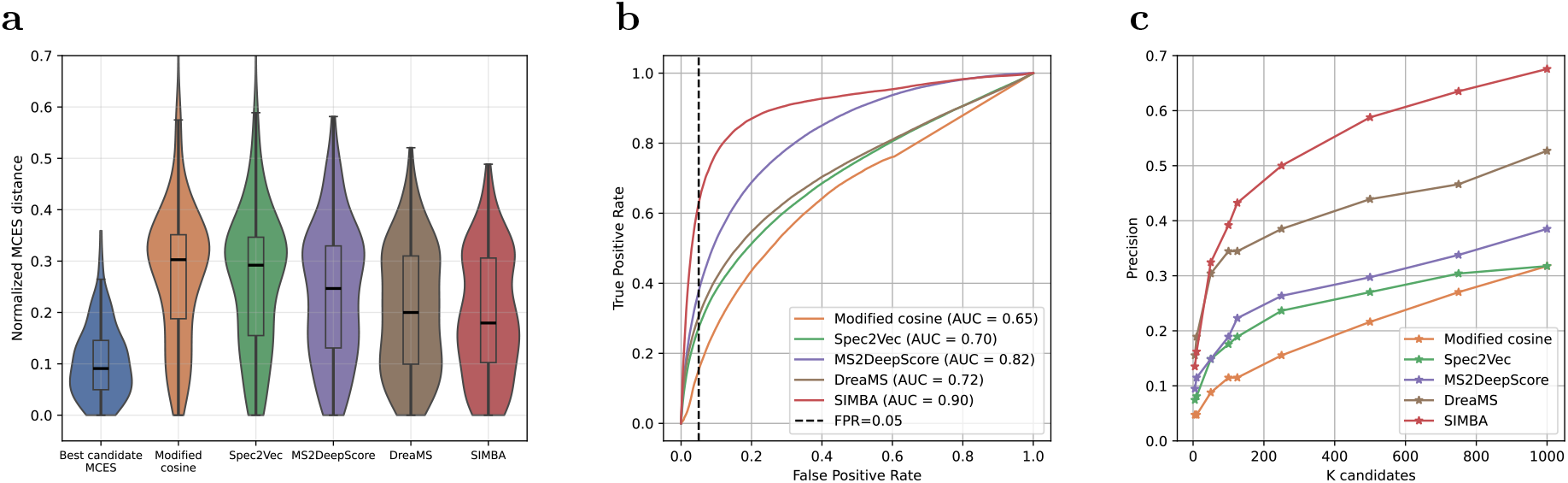
SIMBA outperforms existing methods in analog discovery using CASMI 2022 query spectra against the GNPS community spectral libraries. **(a)** Structural similarity of the top-ranked analog for various similarity measures. The central line indicates the median, the box bounds the interquartile range (25th–75th percentiles), and the whiskers indicate 1.5 times the interquartile range. SIMBA retrieves structurally closer analogs (median normalized MCES 0.18) than modified cosine (median normalized MCES 0.30), Spec2Vec (median normalized MCES 0.29), MS2DeepScore (median normalized MCES 0.25), and DreaMS (median normalized MCES 0.20). **(b)** ROC curves for discriminating true analogs from unrelated molecules, where pairs are binarized using a normalized MCES distance threshold of 0.3. At a false positive rate of 5 %, SIMBA achieves substantially higher true positive rates than competing approaches (SIMBA: 62 %; DreaMS: 30 %; MS2DeepScore: 38 %; Spec2Vec: 28 %; modified cosine: 17 %). **(c)** Ranking performance of analog retrieval. SIMBA more consistently ranks the closest structural analog among the top candidates compared to alternative methods.

**Supplementary Figure 3:**
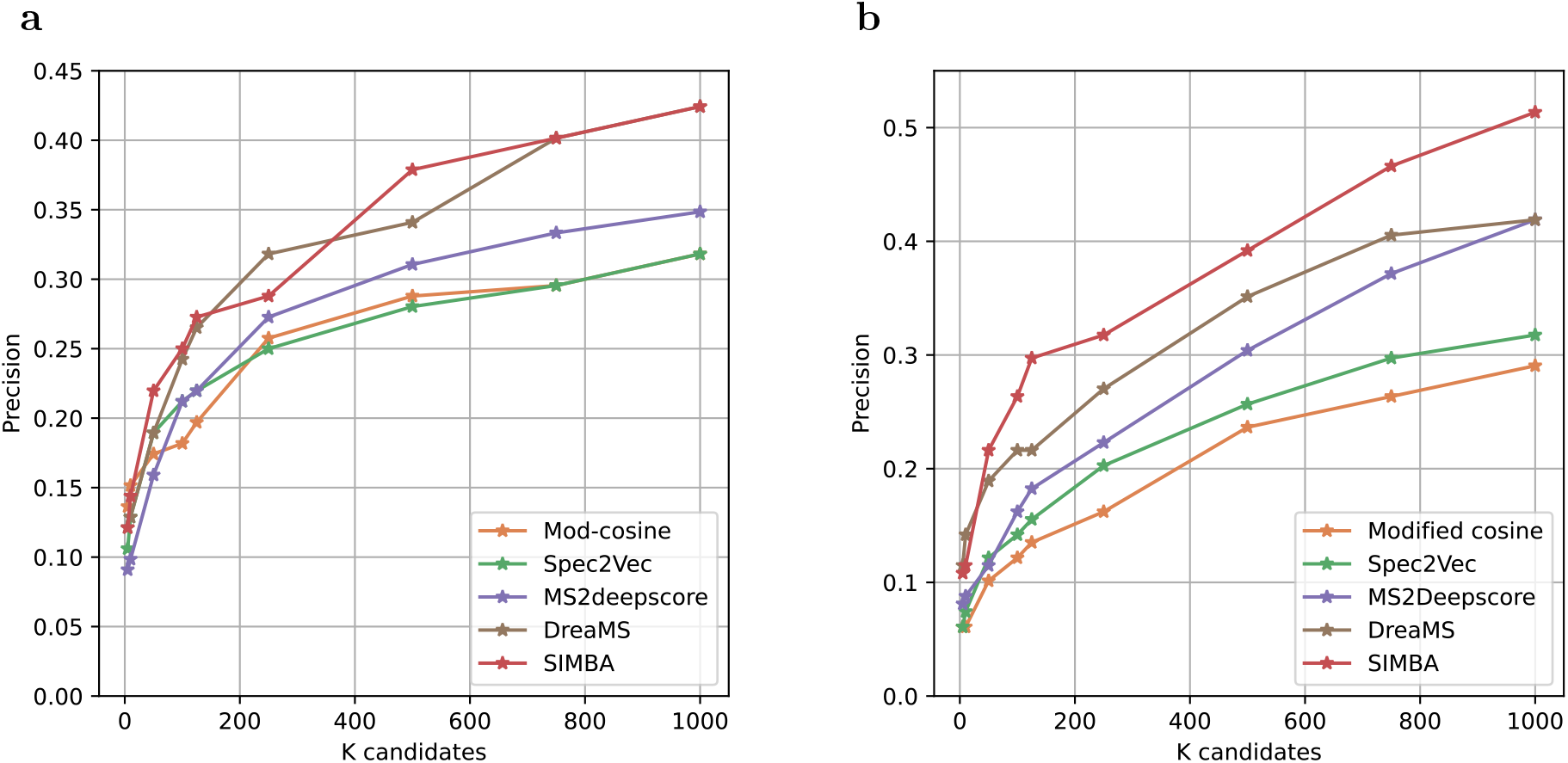
SIMBA improves analog ranking when evaluated using Tanimoto-based structural similarity. **(a)** Ranking of the top candidate based on highest Tanimoto similarity using CASMI 2022 query spectra against the NIST20 + MassSpecGym reference library. **(b)** Equivalent evaluation using the GNPS community spectral libraries. SIMBA more consistently ranks the structurally closest analog among the top candidates compared to alternative methods.

**Supplementary Figure 4:**
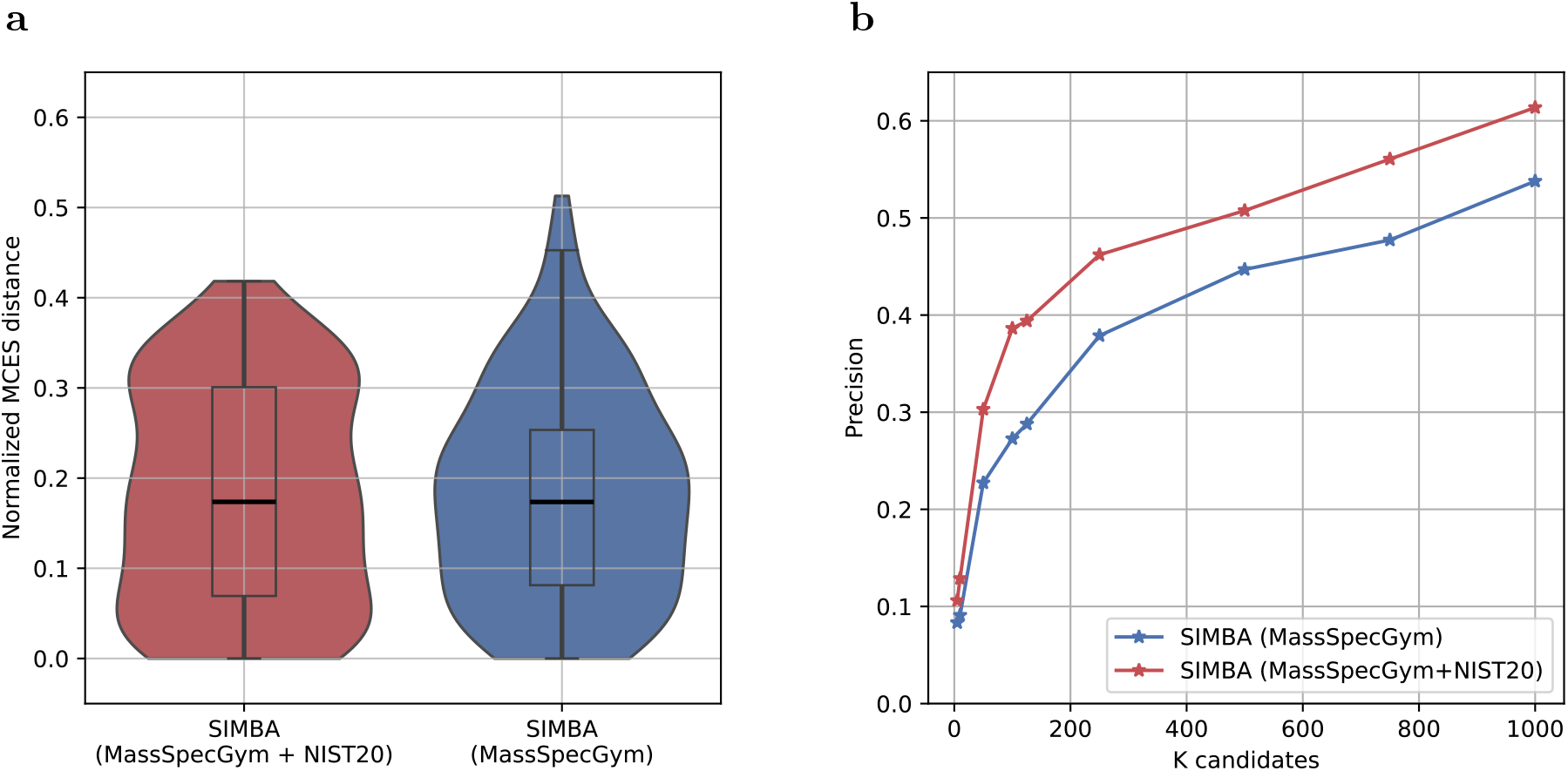
SIMBA trained on public data achieves comparable analog discovery performance to the full model. Performance comparison between a model trained on MassSpecGym and proprietary NIST20 data and a model trained only on MassSpecGym. **(a)** Normalized MCES distance of retrieved analogs for CASMI 2022 queries (lower values indicate higher structural similarity). **(b)** Ranking of the best candidate for CASMI 2022 query spectra.

